# *Woolly* mutation with *Get02* locus overcomes the polygenic nature of trichome-based pest resistance in tomato

**DOI:** 10.1101/2023.10.06.561305

**Authors:** Eloisa Vendemiatti, Inty O. Hernández-De Lira, Roxane Snijders, Tanmayee Torne, Rodrigo Therezan, Gabriela Prants, Carlos Lopez-Ortiz, Umesh Reddy, Petra Bleeker, Craig A. Schenck, Lázaro Eustáquio Pereira Peres, Vagner Augusto Benedito

## Abstract

Type-IV glandular trichomes, which only occur in the juvenile phase of tomato development, produce acylsugars (AS) that broadly protect against arthropod herbivory. Previously, we introgressed the capacity to retain type-IV trichomes in the adult phase from *Solanum galapagense* into the cv. Micro-Tom (MT). The resulting MT-*Get* line contained five loci associated with enhancing the density of type-IV trichomes in adult plants. We genetically dissected MT-*Get* and obtained a sub-line containing only the locus on chromosome 2 (MT- *Get02*). This genotype displayed about half the density of type-IV trichomes compared to the wild progenitor. However, when we stacked the gain-of-function allele of *WOOLLY*, which codes for a HD-ZIP IV transcription factor, MT-*Get02/Wo* exhibited double the number of type-IV trichomes compared to *S. galapagense*. This discovery corroborates previous reports positioning *WOOLLY* as a master regulator of trichome development. AS levels in MT-*Get02/Wo* were comparable to the wild progenitor, although the composition of AS types differed, especially regarding less AS with medium-length acyl chains. Agronomical parameters of MT-*Get02/Wo*, including yield, were comparable to MT. Pest resistance assays showed enhanced protection against whitefly, caterpillar, and the fungus *Septoria lycopersici*. However, resistance levels did not reach that of the wild progenitor, suggesting the specificity of particulars AS types in the pest resistance mechanism. Our findings in trichome-mediated resistance advance the development of robust, naturally resistant tomato varieties, harnessing the potential of natural genetic variation. Moreover, by manipulating only two loci, we achieved exceptional results for a highly complex, polygenic trait, such as herbivory resistance in tomato.

## INTRODUCTION

Despite their minute size, plant trichomes, which are literally the first site of contact with many environmental factors, provide substantial ecological advantages. These hair-like structures help plants tolerate biotic and abiotic stresses by protecting against herbivorous insects and microorganisms, excessive ultraviolet light, heat, drought, and even heavy metals (Hülskamp, 2004; Wagner, 2004; Yang and Ye, 2013; Koul et al., 2021; Zhang et al., 2021). In particular, glandular trichomes are a critical defense mechanism against arthropod herbivores, which warrants the study of their development and biochemistry (Bar and Shtein, 2019).

Insect-plant interactions have been extensively studied, and the tomato (*Solanum lycopersicum*) is among the best-documented model systems (Price et al., 1980; Kennedy, 2003; Wagner, 2004; Prasifka, 2015; Kaur et al., 2023). Trichome-based defenses are critical in several wild tomato species (Kennedy, 2003; Simmons and Gurr, 2005; Maluf et al., 2010; Firdaus et al., 2012; Salinas et al., 2013; Rakha et al., 2017; Vosman et al., 2019; Kortbeek et al., 2021).

Luckwill (1943) classified seven different types of trichomes (I-VII) in the *Solanum* genus, comprising four glandular (I, IV, VI, and VII) and three non-glandular types (II, III, and V) (Fig. 1A). Our work focuses on type-IV glandular trichomes, which are elongated multicellular structures consisting of a single flat basal cell, a short uniseriate two- or three-celled stalk (0.2-0.4 mm), and a spherical gland at the tip (Glas et al., 2012). They differ from the type-V non-glandular trichome only by the presence of the apical gland, and many studies reported a negative correlation between the density of these two trichome types (Fernández-Muñoz et al., 2003; Simmons and Gurr, 2005; Vendemiatti et al., 2017; Vendemiatti et al., 2022).

**Figure 1.**
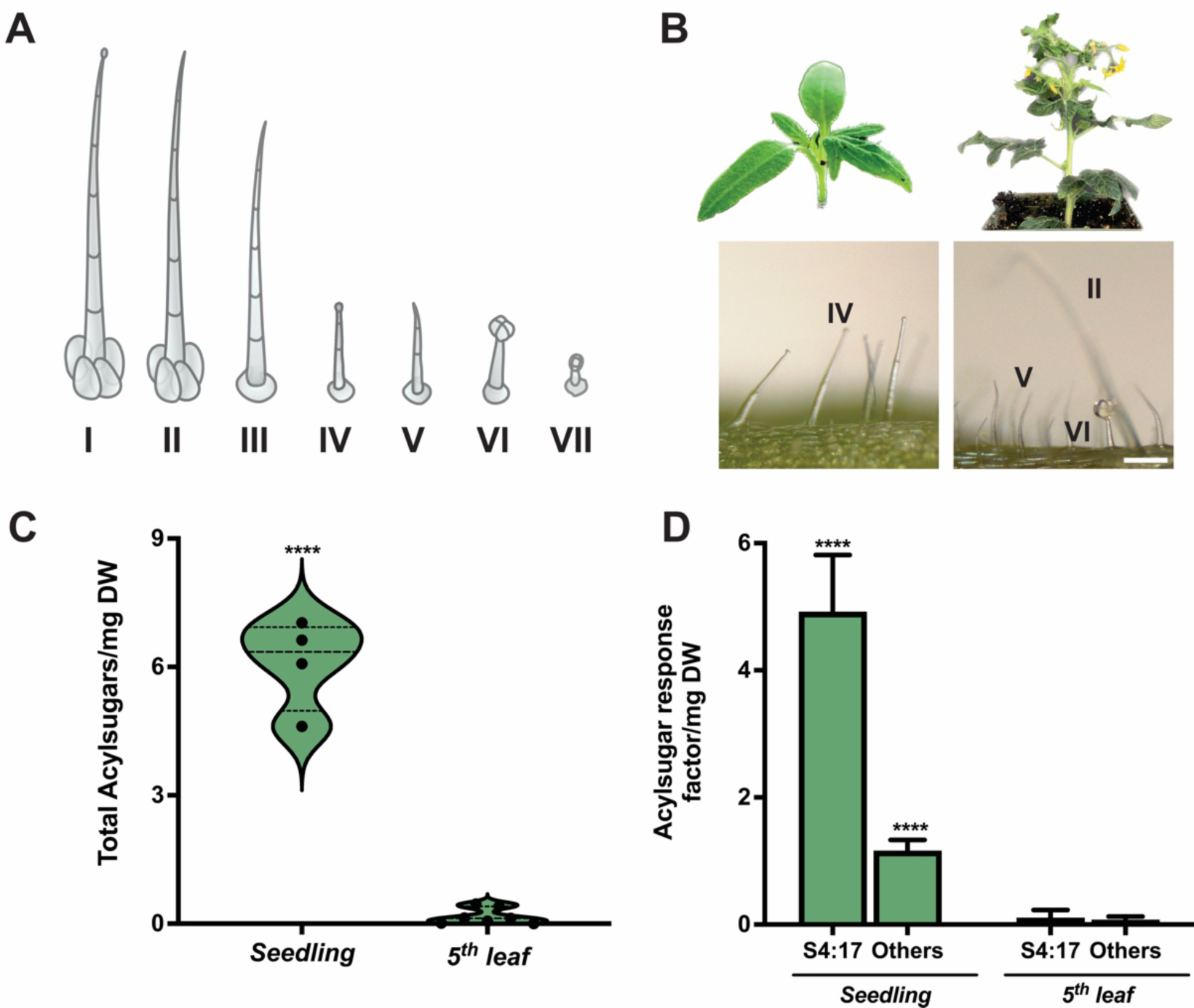
Trichome types and AS profile in the juvenile stage of tomato. **A)** Trichome classification in the *Solanum* genus, according to Luckwill (1943). **B)** Trichome diversity on seedlings and the 5^th^ leaf (adult stage) of *S. lycopersicum* cv. Micro-Tom (MT). **C)** Comparison of total AS concentrations between seedling and adult leaf stage and **D)** Contribution of most prominent AS (S4:17) profile comparison in MT stages. Data are means ± SEM (n≥4 biological replicates). AS nomenclature (S4:17) indicates sucrose core, four total acylations totalizing 17 carbons in those acyl chains of lengths 2, 5, 5, and 5. Asterisks indicate a significant difference between the groups according to the Student’s t-test at *p* < 0.0001. (****). Scale bar = 200 mm.

Type-IV trichomes produce AS, a structurally diverse class of compounds resulting from the acylation of a sugar core (Mutschler et al., 1996; Schilmiller et al., 2008; Leckie et al., 2012; Vosman et al., 2019; Feng et al., 2021; Fiesel et al., 2022). AS are especially effective against arthropod herbivory, including Lepidoptera (*Helicoverpa amigera*, *Spodoptera exigua*, *Phthorimeae operculella*), Acarina (*Tetranychus urticae*), and Hemiptera (*Myzus persicae*) (Goffreda et al., 1990; Hawthorne et al., 1992; Liedl et al., 1995; Simmons and Gurr, 2005; Maluf et al., 2010). AS have also been consistently associated with broad-spectrum resistance against fungal pathogens (Luu et al., 2017). Hence, a trichome-focused approach in breeding for pest-resistant tomato varieties has been pursued for several years, but the optimal genetic combination has yet to be achieved.

Another source of AS includes type-I trichomes, which differ from type-IV structures by their longer stalk length (2 – 3 mm) and globular multicellular base (Luckwill, 1943; Glas et al., 2012; Schilmiller et al., 2012). Type-I trichomes are sparsely distributed across both cultivated and wild tomato species belonging to the *Lycopersicon* and *Lycopersicoides* sections of the *Solanum* genus (Glas et al., 2012; Knapp and Peralta, 2016), rendering them less effective sources of AS. In contrast, type-IV trichomes are persistently abundant during the full lifecycle of some wild tomato species, including *S. habrochaites*, *S. galapagense*, and *S. pennellii*. While type-IV trichomes were previously believed to be absent in cultivated tomatoes (Luckwill, 1943; Simmons and Gurr, 2005), we demonstrated their presence on cotyledons and very young leaves, linking this trait to juvenility (Vendemiatti et al., 2017). In that same study we also noted that the tomato *Woolly* mutant exhibits a high density of type-IV trichomes on its juvenile leaves. *WOOLLY* is a homeodomain leucine zipper IV (HD-ZIP IV) transcription factor, previously described to control type-I trichome formation (Yang et al., 2011). Combined, these findings are consistent with the recent discovery that *WOOLLY* actually controls different trichomes types in a dosage-dependent manner (Wu et al., 2023), and this knowledge opens the possibility for use in molecular breeding.

Aiming to introduce the ability of *S. galapagense* to retain type-IV trichomes in the adult life stage into a cultivated tomato, the tomato introgression line MT-*Galapagos enhanced trichomes* (MT-*Get*) was developed in a cv. Micro-Tom background (Vendemiatti et al., 2022). This MT-*Get* genotype exhibits type-IV trichomes throughout all developmental stages. A mapping-by-sequencing approach revealed that MT-*Get* contains five introgressed loci. Although some individual loci were sufficient for the appearance of type-IV trichomes, the simultaneous presence of the five loci appears to be necessary to increase type-IV trichome density. However, MT-*Get* plants did not exhibit the same level of resistance to whiteflies as observed in *S. galapagense* (Firdaus et al., 2013; Firdaus et al., 2012; Vosman et al., 2019; Vendemiatti et al., 2022). Since both the density of type-IV trichomes as well as the levels of AS in MT-*Get* were inferior to those observed in *S. galapagense*, it was concluded that there are additional genetic factors linked to insect resistance in *S. galapagense*, which is likely to be a complex, polygenic trait.

Considering the quantitative importance of AS and type-IV trichomes in pest resistance, we explored gene interactions to enhance the density of type-IV trichomes in *S. lycopersicum*. To this end, we derived the MT-*Get* sub-line harboring only the chromosome 2 locus from *S. galapagense* that controls type-IV trichome development, called MT-*Get02*. Subsequently, we stacked a mild, non-embryo-lethal *Woolly* allele in homozygosity (*Wo^m^*) (Yang et al., 2011) using the introgression line MT-*Wo^m^* in the same genetic background. Surprisingly, this double variant *Get02/Wo* displayed nearly twice the density of type-IV trichomes on the leaves of adult plants compared to *S. galapagense*. We further explored AS content in these genotypes and assessed resistance against relevant pest and pathogens for tomato cultivation: *Bemisia tabaci* (whiteflies), *Manduca sexta* (caterpillars), and the fungus *Septoria lycopersici* - the etiological agent of *Septoria* leaf spot disease. Our observations showed a compelling epistatic interplay between *Woolly* and the *Get02* locus, critically influencing type-IV trichome development and modulating AS biosynthesis in tomato. Furthermore, our results indicate that a combination of mutations and natural genetic variation may offer a viable solution for overcoming the polygenic nature of pest resistance.

## RESULTS

### 1. AS production in type-IV trichomes present on juvenile leaves of cultivated tomato

Since type-IV trichomes are present on cotyledons of the cultivated tomato, but not on adult leaves (Vendemiatti et al., 2017) (Fig. 1B), we first assessed these glands actively synthesize AS, by profiling these compounds in *S. lycopersicum* cv. Micro-Tom (MT) seedlings. Our data show that MT seedlings indeed produce AS, but that the adult leaves contain only negligible amounts (Fig. 1C). The latter is likely synthesized in type-I trichomes, which are sparsely present on the adult leaves. The most abundant AS to accumulate on MT cotyledons was a tetra-acylated sucrose (S4:17) containing three short chains and an acetyl group (2,5,5,5, see Fig. 1 legend) (Fig. 1C). Interestingly, juvenile leaves exhibit a different AS profile compared to leaves developed at the adult plant stage. While AS with a 12-carbon long acyl chain were present in cotyledons, this type was absent in later leaves (Supplemental Fig. S3).

### 2. Type-IV trichomes density in the Get02/Wo double variant tomato line

Previously, we identified five chromosomal segments in the introgression line MT-*Get* inherited from the wild progenitor *S. galapagense* (LA1401), associated with persistent development of type-IV trichomes throughout plant life. One of the sub-lines resulting from this genetic dissection of MT-*Get* contained only one of these segments and was named MT-*Get02* (Fig. 2A and B), as this is the locus on the long arm of chromosome 2 of *S. galapagense* between coordinates 51,924,000 – 52,413,200 (genome assembly SL4.0). This locus contains 203 annotated genes and was previously associated with a QTL for resistance to whitefly (Firdaus et al., 2012; Firdaus et al., 2013).

**Figure 2.**
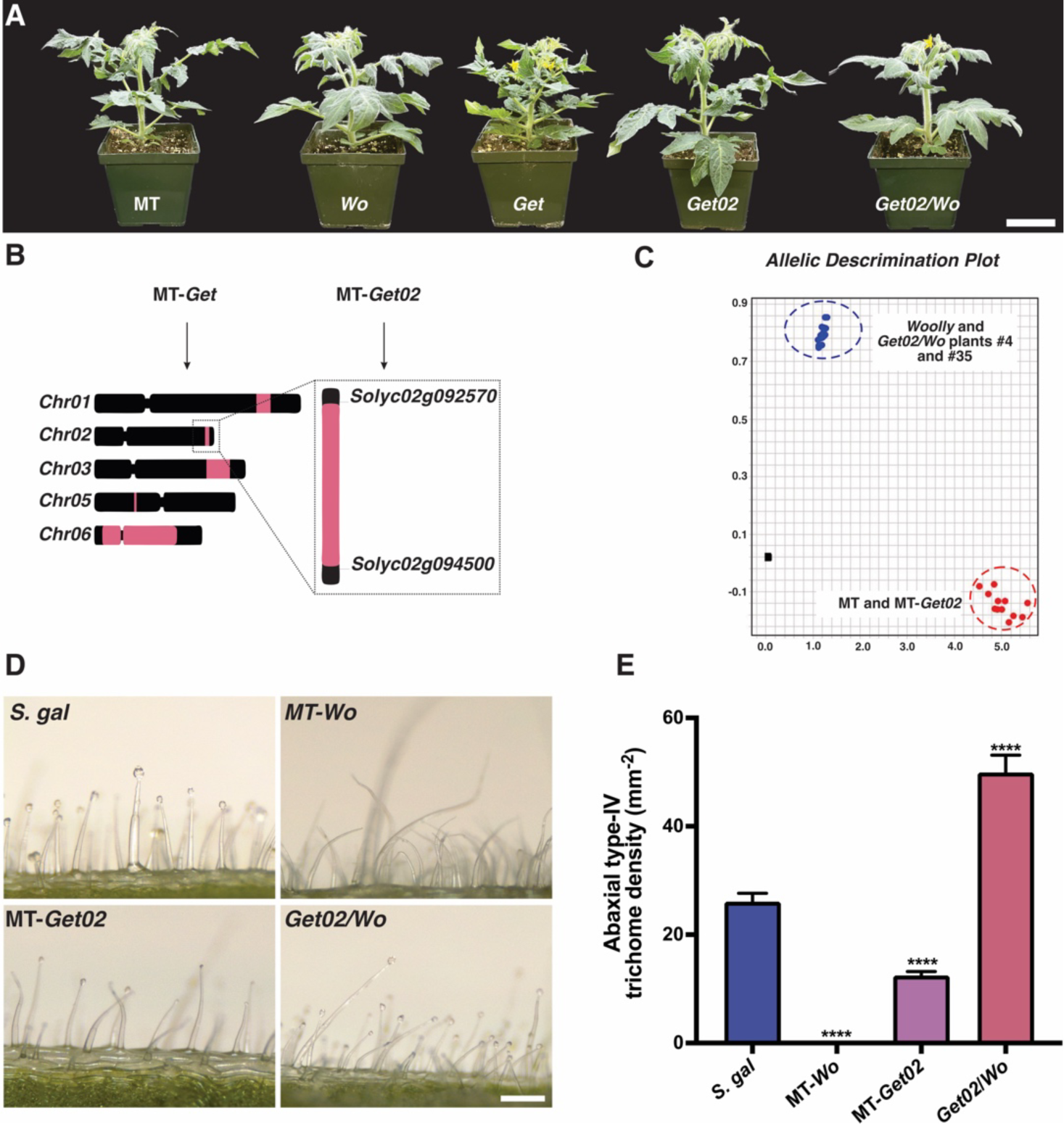
Phenotypic characterization of tomato introgression lines in the cv. Micro-Tom background. **A)** Comparison of *S. lycopersicum* cv. Micro-Tom (MT), MT-*Woolly* (*Wo*), MT-*Galapagos enhanced trichomes* (MT-*Get*), MT-*Galapagos enhanced trichomes02* (MT-*Get02*), and the double variant *Get02/Wo.* Scale bar = 5 cm. **B)** Approximate locations of the introgressions in MT-*Get* and MT-*Get02* chromosomes. The segments highlighted in pink are related to the *S. galapagense* fragments in the MT genome. **C)** PACE analysis results show that MT-*Get02* plants #4 and #35 have the *Wo^m^* gain-of-function mutation in homozygosis. **D)** Representative leaf abaxial surface images show the presence/absence of type-IV trichomes. Scale bar = 200 mm. **E)** Quantification of type-IV trichomes in the tomato lines used in this work. Data are the means of 50 biological replicates ± SEM. Asterisks indicate significant differences compared to *S. galapagense* (*S. gal*), according to Dunnett’s test at *p* < 0.0001. (****).

*Wo^m^* is a mild gain-of-function mutation in the gene *WOOLLY* (*Solyc02g080260*) that increases the density of digitate trichomes (mainly types III and IV, Vendemiatti et al., 2017) and results in a hairy phenotype (Fig. 2 - Yang et al., 2011; Wu et al., 2023). This gene is located 9.16 Mb upstream of the MT-*Get02* locus. To investigate the combined effect of the *Get02* locus and *Wo* mutation on the type-IV trichome pathway, we sought to produce a double variant line; combining MT-*Get02* with a near isogenic line (NIL) in the cv. MT background that contains the mild *Wo^m^* mutation originating from the accession LA0715 (*S. lycopersicum*). While strong *Wo* alleles are lethal to embryos in a homozygous state (Yang et al., 2011), the *Wo^m^* allele is also a spontaneous, viable mutation in the cv. Rutgers (LA0258).

To obtain the double Get02/Wo line, a F2 segregating population resulting from the crossing between MT-*Get02* and MT-*Wo^m^* was screened for type-IV trichome density (Supplemental Fig. S1). Out of 98 plants screened, 18.37% exhibited a significant increase in type-IV trichomes compared to the parental MT-*Get02* (Supplemental Fig. S1). This observation fits with the expected Mendelian proportion of 3/16 (18.75%) for independent segregation of two genes, where one contains homozygous alleles (*Get02*) and the other has homozygous/heterozygous alleles (*Wo^m^*). Alternatively, this proportion could indicate that the two genes are not segregating independently which is a possibility, seen that both are located on chromosome 2, within less than 50 cM.

Among the F2 plants identified to exhibit high densities of type-IV trichomes, one plant (line #4 in Supplemental Fig. S1) developed nearly twice the number of trichomes compared to *S. galapagense* (Fig. 2D and E). Interestingly, this plant was sticky to the touch, an indicative for the presence of AS. Besides plant #4, we also selected plant #35 and confirmed both contained the two loci, *Get02* and *Wo^m^*. Self-fertilization of plant #4 and #35 produced F3 seeds, whose seedlings were subsequently screened for the presence of both loci in homozygosity. The presence of the homozygous *Get02* alleles was confirmed using CAPS markers and the *Wo^m^* homozygous alleles by PACE analysis (Fig. 2C), thus establishing the *Get02/Wo* double variant lines. Next, we analyzed the AS profile in homozygous plants derived from the double *Get02/Wo* lines and tested the performance of this material in herbivory assays with whiteflies and caterpillars, and a fungal disease exposure.

### 3. Get02/Wo acylsugars profiling

Due to the stickiness of *Get02/Wo* leaves, suggestive of high AS production, we profiled this lineage by both liquid- and gas chromatography-mass spectrometry (LC-MS and GC-MS) (Fig. 3, Supplemental Fig. S2 and Fig. S3). *Get02/Wo* produced similar amounts of total AS as *m/z* peaks in *S. galapagense* (Fig. 3B). However, the AS composition of *Get02/Wo* contains mainly tetra- acylated sucrose (S4:17, m/z 681.3, Fig. 3A and C). Compared to *S. galapagense*, *Get02/Wo* produces significantly lower concentrations of S3:22(5,5,12) and S4:24(2,5,5,12) structures.

**Figure 3.**
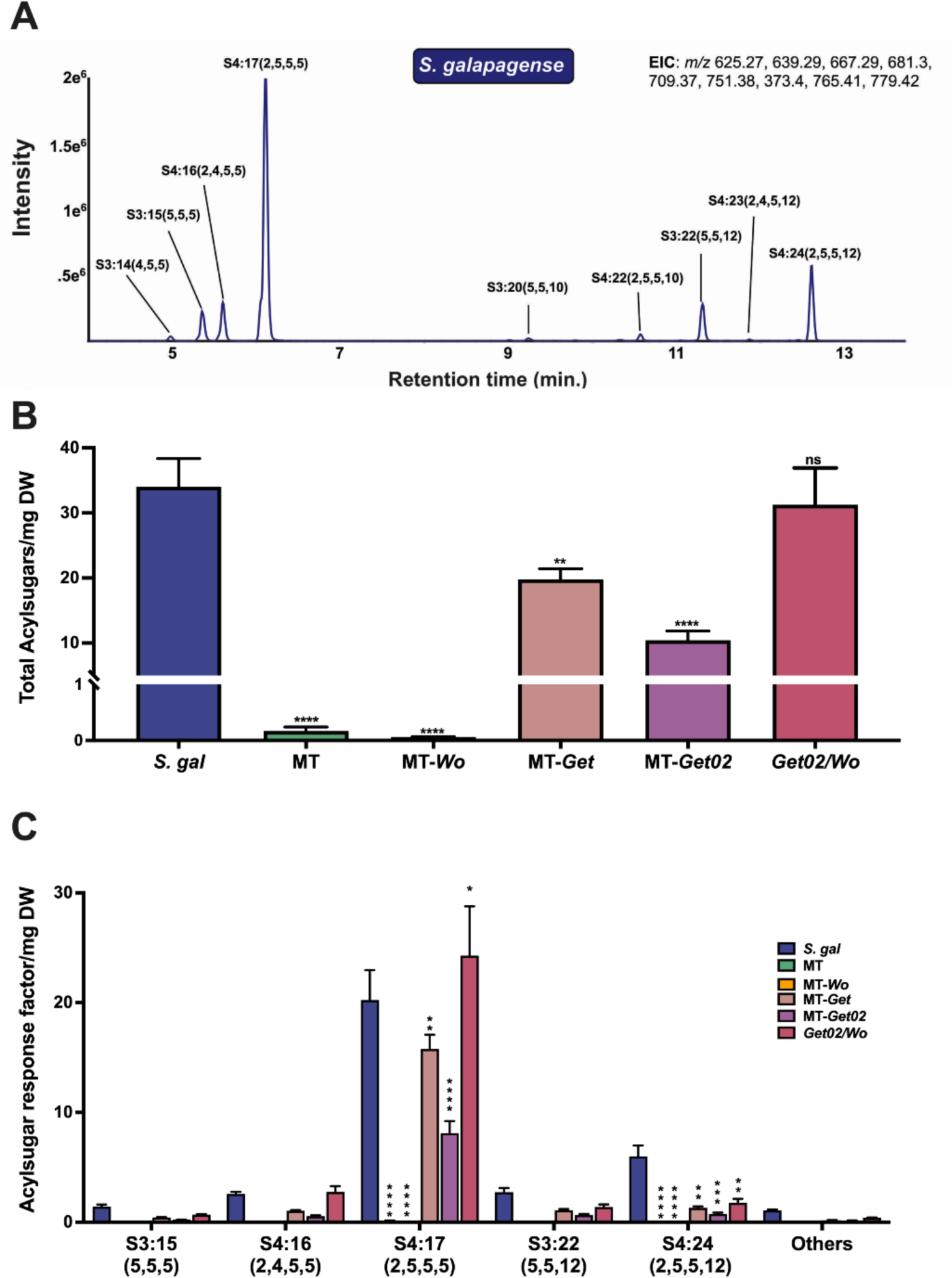
AS profiles in the study material. **A)** AS profiles of leaf extracts of *S. galapagense* and its derivative introgression lines in cv. MT. **B)** Total AS quantification across the introgression lines. Individual AS were quantified based on the peak area of the formate adduct and corrected for the internal standard (telmisartan) and the dry leaf weight. **C)** Quantification of the main individual AS types (A) was added, resulting in the total AS in each line. Data are the means of ≥5 biological replicates ± SEM. Asterisks indicate significant differences compared with *S. galapagense*, according to Dunnett’s test at *p* < 0.01(**), *p* < 0.001 (***), and *p* < 0.0001. (****).

Furthermore, GC-MS analysis was further performed to evaluate the composition of acyl chains in each genotype. Our results revealed that the main difference in mature leaves of the cultivated tomato (MT) compared to *S. galapagense* is the lack of AS with medium acyl chains (10-12 carbons in length). However, AS containing 12 carbon long acyl chains are produced in all introgression lines with type-IV trichomes and, to a lesser extent, in MT seedlings (Supplemental Fig. S3).

### 4. Pest Resistance Assessment in Get02/Wo

To investigate the biological effect of the altered AS profile in our material, we performed bioassays with relevant biotic interactors of tomato. We conducted two no-choice assays to evaluate the survival and oviposition rates of the pest insect *Bemisia tabaci* among the genotypes (Fig. 4A, Supplemental Fig. S4). As expected, significantly fewer eggs were deposited on *S. galapagense* compared to MT, and the single-locus introgressions of MT-*Wo* and MT-*Get* did not result in altered oviposition rates. A slight reduction in oviposition was observed in the double variant *Get02/Wo,* but this difference was not significant when compared to *S. galapagene* (Fig. 4A). However, significantly fewer eggs were laid on MT-*Get02/Wo* when compared to MT in Petri dish assays (Supplemental Fig. S4). In the same experiment, there was no significant difference in the survival rate among the genotypes. These results suggest that MT-*Get02/Wo* can provide resistance to whitefly infestation by deterring oviposition.

**Figure 4.**
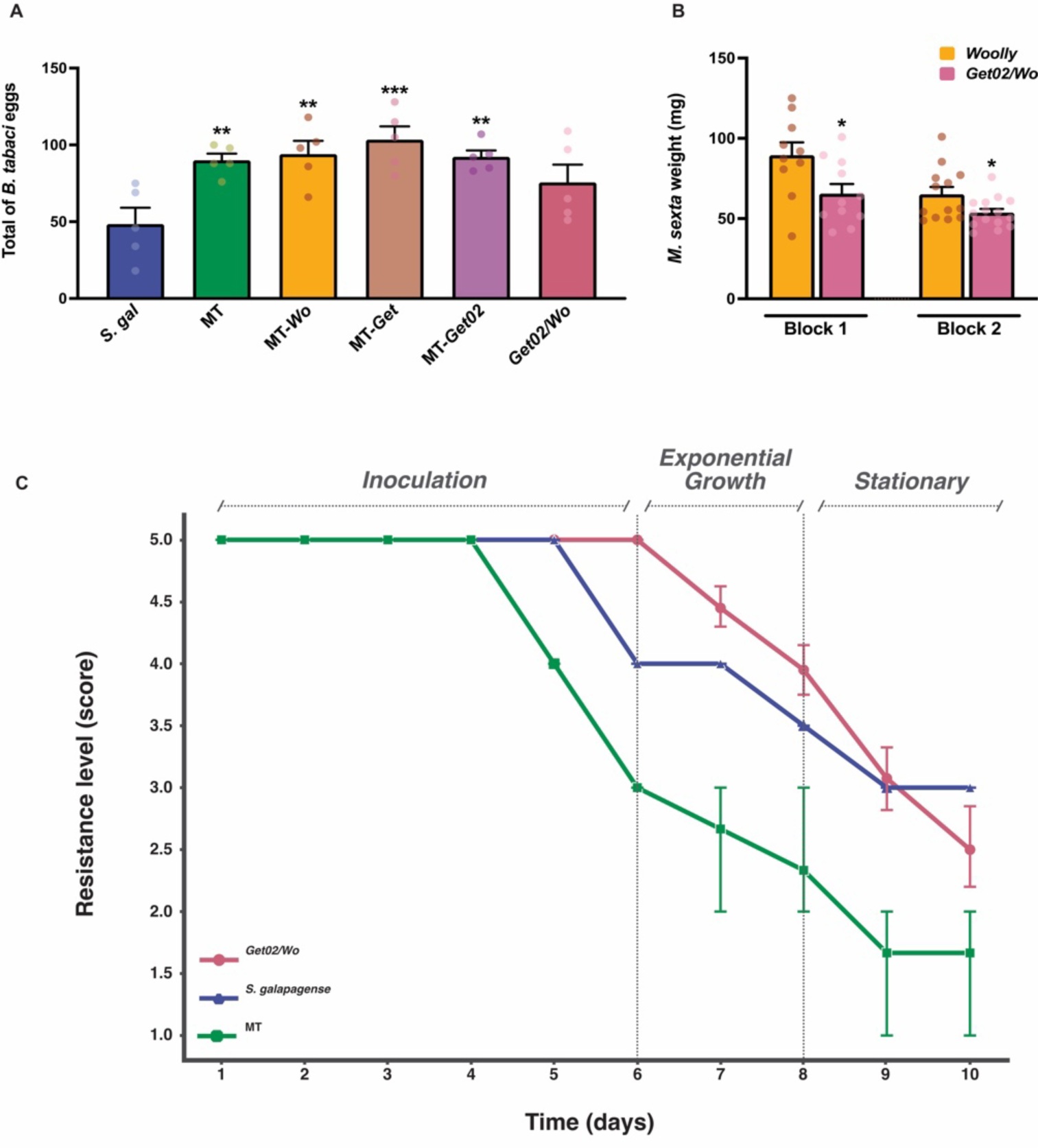
Insect Resistance Assessment. **A)** Total number of *B. tabaci* eggs, two days post infestation on leaves of different genotypes using clip cages. Data are means of five biological replicates ± SEM. *p* < 0.01 (**), *p* < 0.001 (***) according to Dunnett’s t-test compared to *Solanum galapagense* (*S. gal*). **B)** Weight of *M. sexta* **larvae** feeding on MT-*Wo* and *Get02/Wo* for 7 days. Each block represent an experiment repetition. **C)** Infestation score of *Septoria* Leaf Spot (SLS) in MT, *S. galapagense* and *Get02/Wo* plants during 10 days of infestation.

We also evaluated the effect of increasing plant AS on the development of *Manduca sexta* caterpillars when they were fed MT-*Wo* compared to *Get02/Wo* leaves. Due to experimental limitations, only two genotypes could be analyzed. Thus, we decide to compare the MT-*Wo,* which has a high density of non-glandular trichomes with very low amount of AS (Fig. 3B), to *Get02/Wo*, the genotype with a high density of type-IV trichomes accumulating high concentrations of AS. After 7 days, the *Get02/Wo* fed caterpillars were significantly smaller than those fed MT-*Wo* leaves (Fig. 4B). These results suggest that the amount and the AS profile in *Get02*/*Wo* can delay the growth of caterpillars.

AS have also been shown to contribute to pathogen resistance, based on a lower spore germination and smaller necrotic lesions (Luu et al., 2017). Thus, we tested the resistance of the *Get02/Wo* to *Septoria* Leaf Spot (SLS) infection, which is a well-establish study system in our laboratory. Our evaluation showed that the wild species *S. galapagense* has a medium-high tolerance (3.0/5 score) to this fungus and the incubation period of the fungus was delayed by one day compared to the MT control (Fig. 4C). While on the *Get02/Wo* plants the incubation time was delayed by two days, and plants displayed prolonged tolerance throughout the pathogen exponential growth phase, compared to the wild species. However, *Get02/Wo* plants were unable to maintain this resistance throughout the entire infection period. With these results, we conclude that the *Get02/Wo* lineage has a medium resistance (2.5/5) for this disease and that the new genotype improved the tolerance level to SLS compared to the cv. MT (1.6/5 score).

### 5. Agronomical Parameters in Get02/Wo

The *Get02/Wo* parental line, MT-*Get*, exhibits several traits that could affect agronomical performance, such as reduced fruit weight (Vendemiatti et al., 2022). Therefore, we evaluated *Get02/Wo* performance also because there is a concern that breeding based on high trichome densities and enhanced specialized metabolism may negatively impact yield due to the deviation of energy from primary metabolism that contributes to fruit size and, partially, to taste (Tieman et al., 2017). *Get02/Wo* plants have the same height and flowering time as MT (Supplemental Fig. S5A and B). In contrast, MT-*Get* plants are smaller and take longer to flower than MT and *Get02/Wo* (Supplemental Fig. S5A and B; see also Fig. 2A). There was no statistical difference in stem diameter among the lines (Supplemental Fig. S5C).

MT-*Get* and MT-*Get02* produced, respectively, the lowest total fruit weight and the lowest fruit Brix, compared to all other genotypes analyzed (Fig. 5). Interestingly, MT-*Wo^m^* showed the highest Brix among the genotypes evaluated. Our double variant line *Get02/Wo* did not show significant differences in total fruit weight and Brix, and this way, it is very similar to the control MT (Fig. 5).

**Figure 5.**
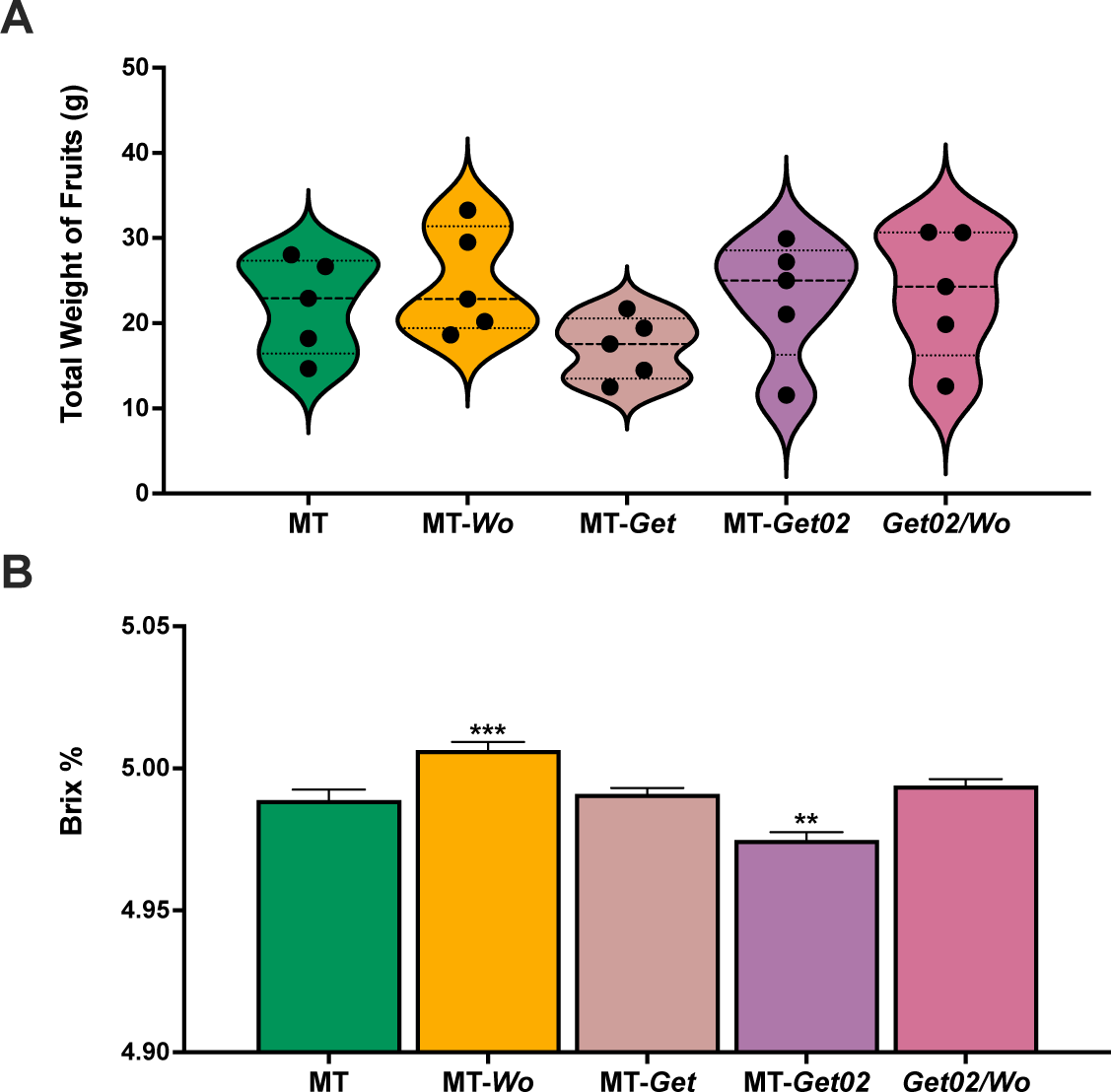
Impact of genetic introgressions on yield parameters. **A)** Total weight of fruits (g) produced by each genotype. Data are means of five biological replicates ± SEM. According to Dunnett’s t-test and using MT as a control, the data is not significantly different at *p* < 0.05. **B)** Soluble solids content (% Brix) for each genotype. Data are the mean of ten fruits per genotype ± SEM. *p* < 0.01 (**), *p* < 0.001 (***) according to Dunnett’s t-test using MT as control.

## DISCUSSION

### The Get02 natural genetic variation enables the occurrence of type-IV trichomes in the adult phase of tomato development

In 2005, Simmons and Gurr proposed a trichome-based resistance to minimize pesticide use by introducing trichome traits from wild accessions into cultivated tomatoes. Such an approach is grounded in the understanding that glandular trichomes and their specialized metabolites contribute to pest and pathogen resistance (Weinhold and Baldwin, 2011; Firdaus et al., 2013; Rakha et al., 2017; Luu et al., 2017; Feng et al., 2021). While numerous studies have delved into this field, a commercial tomato line with trichome-based resistance has yet to be achieved. Basically, the main forms of resistance are based on metabolites produced in type-IV and type-VI trichomes. In terms of pest resistance deriving from type-IV trichomes—the most predominant type of acylsugar-producing trichomes in some wild tomato species—the lack of these structures in adult plants of cultivated tomato accessions seems to be the main obstacle in developing plants with broad, enduring resistance against arthropod herbivores.

In a previous study, we have shown that cultivated tomatoes display type-IV trichomes in the juvenile phase of plant development (Vendemiatti et al., 2017). Herein, we confirm that they are metabolically active and accumulate AS (Fig. 1C and Supplemental Fig. S3). However, the intricate genetics dictating their persistence in the adult phase of the plant could be the obstacle preventing the success in obtaining a commercial tomato line with trichome-based pest resistance (Vendemiatti et al., 2022; Mutschler et al., 2023). Understanding the key genetic factors that could stabilize type-IV trichome development in the cultivated tomato prompted us to transfer this trait from *S. galapagense* to the tomato model system Micro-Tom (MT), creating the multi-locus MT-*Get* lineage (Vendemiatti et al., 2022). In the present work, we advanced the dissection of the MT-*Get* into derivative sublines, each with a single locus from the wild progenitor. We initiated this exploration with MT-*Get02*, the subline that carries the fragment from chromosome 2 from *S. galapagense*. This decision was based on the preliminary data from Vendemiatti et al. (2022) that attributed this locus as critical for the persistent development of type-IV trichomes in MT-*Get*. Further reinforcing our choice, we identified a correlation with insect resistance in previous studies pointing to this same genomic region by i) updating the coordinates on chromosome 2 from the QTLs outlined by Salinas et al. (2012) and Smeda et al. (2016) using the latest version of the S*. lycopersicum* cv. Heinz genome assembly (SL4.0), and ii) comparing these ranges with the introgression obtained by Vendemiatti et al. (2022). We are currently using this approach to identify the culprit gene on chromosome 2 involved with trichome development and pest resistance.

Since type-IV trichomes are present in the mature phase of different tomato wild species, it is plausible that such species employ a unique set of genes, potentially a genetic network, influencing the presence of type-IV trichomes at a specific stage of the plant lifecycle (Maliepaard et al., 1995; Momotaz et al., 2010; Leckie et al., 2012; Firdaus et al., 2013; Smeda et al., 2016; Mata-Nicolás et al., 2021; Vendemiatti et al., 2022). Interestingly, the genomic region encompassed by *Get02* has been associated with type-IV trichome development in three wild species of tomato: *S. habrochaites*, *S. pennellii,* and some accessions of *S. pimpinellifolium* (Maliepaard et al., 1995; Momotaz et al., 2010; Salinas et al., 2013; Smeda et al., 2016). Altogether, these pieces of information suggest the existence of a locus on chromosome 2 driving the type-IV trichome developmental pathway in the *Solanum* genus.

### The Woolly mutation plays a pivotal role in increasing the density of type-IV trichomes in the adult phase, when combined with the Get02 locus

The MT-*Get02* subline mirrored the MT overall appearance (Fig. 2A), except for the persistence of type-IV trichomes in the adult phase. Notably, unlike MT-*Get*, containing 5 introgression sections, this subline has no yield penalties (Fig. 5A). To enhance the density of type-IV trichome in MT-*Get02*, we examined its interaction with *Woolly* (*Wo*), a spontaneous tomato mutation initially linked with type-I trichome formation (Yang et al., 2011) and further characterized as having a high density of type-IV trichomes in the juvenile phase (Vendemiatti et al., 2017). The increased density of type-IV trichome in the adult phase of *Get02/Wo* supports the notion that *Wo* can enhance type-IV trichome formation (Vendemiatti et al., 2017), and it suggests that *Get* loci play a role in controlling juvenility (Vendemiatti et al., 2022). Interestingly, recent research has shown that the *Woolly* mutant can increase the density of all digitate trichome types (Wu et al., 2023), including types I, II, III, IV and V. While types I, II and III are sparsely present in all phases of tomato development, types IV and V predominate in the juvenile and in the adult phases, respectively, with their densities showing a negative correlation throughout tomato development (Vendemiatti et al., 2017). Since type-IV and type-V trichomes differ morphologically solely by the presence of a single-cell gland at the tip of the former, they likely share the same initial developmental pathway. Therefore, determining the genetic identity of *Get02* may contribute to our understanding of the genetic network downstream of *WOOLLY* in specifying different digitate trichome types (Wu et al., 2023). According to the model proposed by Wu et al. (2023), elevated *WOOLLY* levels result in digitate trichome differentiation. In contrast, low levels lead to the formation of peltate trichomes (types VI and VII). This phenomenon is also evident in *Get02/Wo*, which exhibits a reduced density of peltate (mainly type-VI) trichomes (Supplemental Fig.S6). Type-VI trichomes also produce natural insecticides, such as plastid-derived sesquiterpenes and methyl ketones, which are absent in cultivated tomatoes but present in the wild species *S. habrochaites* and *S. habrochaites* f. *glabratum* (Sallaud et al., 2009; Therezan et al., 2021). These findings suggest that it may be challenging to engineer both type-IV and type-VI trichome-based pest resistance in the same tomato genotype. We also observed the presence of branched trichomes in MT-*Wo^m^* as well as *Get02/Wo* (Supplemental Fig. S6), consistent with our previous findings (Vendemiatti et al., 2017). This resembles the trichomes present in *Arabidopsis thaliana*, where branching is associated with the extent of DNA endoreduplication (Schwab et al., 2000).

### The Get02/Wo genotype produces elevated levels of AS, with a molecular profile distinct from that of the wild parental S. galapagense but capable of imparting pest resistance

The higher density of type-IV trichomes in *Get02/Wo*, which surpasses the trichome density of the wild progenitor, is likely the reason behind its increased production of AS (Fig. 3B and C). However, the composition of individual AS structural variants differed considerably. Notably, *Get02/Wo* synthesized significantly more S4:17 than *S. galapagense* (Fig. 3C). This AS is the predominant type in *S. lycopersicum* cv. M82 (Kim et al., 2012; Schilmiller et al., 2015; Fan et al., 2016) and, interestingly, it is also the most abundant type in *Get02/Wo*. An important question that arises from this is the effectiveness of these AS types for pest resistance. Testing this hypothesis has proven challenging because cultivated tomatoes, such as M82, only produce minimal amounts of AS, likely attributable to the absence of type-IV trichomes and the sparce density of type-I trichomes.

We conducted pest evaluation performance to assess the impact of the *Get02/Wo* trichome and AS profile on two insect orders with distinct feeding mechanisms: whiteflies (*Bemisia tabaci*) and caterpillars (*Manduca sexta*). This choice was based on the literature that shows AS from the Solanaceae family acting negatively on these orders (Weinhold and Baldwin, 2011; Firdaus et al., 2013; Vosman et al., 2018; Wang et al., 2022a). Our results showed that *Get02/Wo* plants had an adverse impact on whiteflies (less oviposition than the control MT) and caterpillars (individuals were smaller in the AS presence than when fed with control leaves) (Fig. 4). Regarding the disease resistance evaluation, the *Get02/Wo* plants appear to show improved tolerance against SLS. We noticed an incubation delay of two days and an elevated resistance level during the two crucial days of exponential fungal growth. This can be a good starting point in commercial cultivation since it would provide farmers more time to act with measures to contain this disease, either by using compounds approved for organic cropping (*e.g*., copper) or synthetic fungicides (e.g., azoxystrobin, difenoconazole, dithiocarbamate).

The pest evaluation panel showed that the trichome/AS profile combination in *Get02*/*Wo* plants currently does not offer the same level of resistance against insects and fungi as that achieved in the parental *S. galapagense*. Since these plants produce the same total amount of acylsugars while sporting double the density of type-IV trichome compared to the parental wild species (Fig. 2B), the bottleneck is likely to be the specific composition of the AS mélange that the plants are accumulating. Acyl chains with a length range of C6-C9 are commonly correlated to caterpillar’s acylsugar-based resistance (Van Dam and Hare, 1998; Luu et al., 2017; Feng et al., 2021; Wang et al., 2022). Recently, the AS S3:15 and S3:21 were predicted to provide whitefly resistance (Kortbeek et al., 2021). Additionally, Puterka et al. (2003) established that octanoate and decanoate fatty acid chains had the highest broad-spectrum insecticidal activity. Based on our results, *S. galapagense* produces AS using acyl chains with 2, 4, 5, 10, and 12 carbons, so it does not accumulate octanoate acyl chains (Fig. 3). *Get02/Wo* has the same production of S3:15 as the wild species. Therefore, we hypothesize that introgressing the genetic components responsible for longer acyl chains (10 or 12 carbons) might be the key to further enhancing the insect resistance of *Get02/Wo* plants.

### Concluding remarks and perspectives

In conclusion, we have engineered a novel tomato genotype combining a specific mutation affecting trichome differentiation fate and a natural genetic variation likely affecting heterochronic alterations in juvenile/adult phase. The *Get02/Wo* exhibits enhanced resistance to insects and a fungal disease, overcoming the polygenic nature normally associated with pest resistance. Since *Get02/Wo* had yield and fruit Brix levels comparable to those observed in MT (see Fig. 5), this allelic combination is a compelling candidate for further evaluation in commercial tomato cultivars and hybrids within pre-breeding programs. Next, we aim to further refine the acylsugar composition to bolster its protective properties and achieve resistance levels on par with *S. galapagense* and other wild accessions. We postulate that increasing the accumulation of long acyl chains containing AS could achieve this goal. Consequently, our future endeavors will focus on genes related to branched-chain amino acid metabolism, which provide acyl chain precursors and trichome-specific fatty acid synthases that catalyze acyl chain elongation to yield *n*-decanoate (10C) and *n*-dodecanote (12C) acyl chains (Slocombe et al., 2008; Ji et al., 2023). Moreover, the forthcoming discovery of *Get02* genetic identity will significantly enhance our comprehension of the developmental pathways regulating the differentiation of each type of digitate trichomes.

## MATERIAL AND METHODS

### 1. Plant Material

Seeds of the tomato (*Solanum lycopersicum*) varieties and the natural variations in the cv. Micro-Tom (MT) background were from the University of São Paulo (Piracicaba, Brazil) repository (www.esalq.usp.br/tomato). The MT seeds were originally obtained from Dr. Avram Levy (Weizmann Institute of Science, Israel) in 1998 and subsequently maintained (through self-pollination) as a true-to-type cultivar. The introgression process of natural variation into the MT cultivar was previously described (Pino et al., 2010; Carvalho et al., 2011; Lombardi-Crestana et al., 2012; Vendemiatti et al., 2022). Plants were cultivated in a growth chamber with a temperature of 28°C, 13h photoperiod, 60% ambient relative humidity, and 150–250 μmol photons m^−1^ s^−2^ PAR irradiance.

### 2. Double variant lineage <u>Get02/Wo</u>

First, we segregated each of the introgressed *S. galapagense*’s chromosomal segments (Vendemiatti et al., 2022) into distinct sublines of MT-*Get* using CAPS markers to enable evaluating the impact of each locus on the trait “density of type-IV trichomes on the 5^th^ leaf”. For that, a segregating population (BC7F2) of MT-*Get* containing 315 plants was screened for the presence of type-IV trichomes on leaves developed during the mature stage of the plant. Selected selfed plants were screened repeatedly until isolines with a single chromosomal segment from the wild progenitor were obtained. With that, each derivative subline from MT-*Get* contains only one homozygous fragment from the wild progenitor, *S. galapagense* (LA1401). Supplemental Table S1 lists the primers used in this work. One of the lineages generated by this process is MT- *Get02,* which is described here. To produce the double variant line, *Get02/Wo*, MT-*Get02* emasculated flowers received MT-*Wo^m^*pollen, and the F1 plants were self-fertilized to produce the F2 generation, which was screened for type-IV trichome quantification (Suppl. Figure S1).

### 3. Trichome Morphometrics Analysis

The trichome quantification was carried out using the methodology described by Vendemiatti et al., (2017, 2022). In summary, clear nail polish fixed leaflet samples in glass slides. A Styrofoam block supported the slide so we can photograph the trichomes in the lateral view, allowing for precise classification and quantification of each type. A Leica Stereoscope EZ4W (Wetzlar, Germany) set to 55x magnification was used to photograph the trichomes. Trichome densities were calculated as trichome counts per unit leaf area.

### 4. PACE-based SNP genotyping of Wo^m^ mutation

Since developing a CAPS marker for the *Wo^m^* allele was cumbersome, we decided on employing PACE. A SNP of interest was selected to develop the PCR allelic competitive extension (PACE) marker, and allele-specific primers were designed with a common reverse primer (Integrated DNA Technologies). The PACE PCR components included 5 µL of PACE Genotyping Master Mix (2X) (3CR Bioscience, Essex, UK) containing FAM, HEX, and ROX fluorophores, 0.11 µL primer assay mix (72X), 2 µL template DNA, and 3 µL of molecular biology grade water. The PCR conditions were adopted in three stages as follows: (1) one cycle at 94 ℃ for 15 min, (2) 10 cycles each including template denaturation (94℃, 20s) and annealing/extension with a drop of 0.8℃ per cycle (65 to 57℃, 60s), (3) 30 cycles each comprising denaturation at 94℃ for 20s, and annealing/extension at 57℃, for 60s (3crBioscience, UK). Further, 3 to 9 cycles of final denaturation and annealing/extension were performed to increase the amplification and generate dense, well-separated clusters. Genotypes were clustered using StepOne plus auto caller (Applied Biosystems). The primer sequences used in this analysis are described in the Supplemental Table S1.

### 5. Acylsugar Profiling

AS were separated and detected using LC-MS in negative ion mode following extraction from leaf dips. Individual AS are labeled according to their annotated structure - e.g., S3:14(4,5,5) indicates an acylsugar with a sucrose core and three acyl chains totaling 14 carbons of lengths 4, 5, and 5 carbons in each acyl chain. AS were extracted from a single fully expanded leaf from 5- week-old plants, which represents the adult phase in MT (Vendemiatti et al., 2017). A leaf was placed into 1 mL of extraction solvent (3:3:2 acetonitrile:isopropanol:water + 0.1% [v/v] formic acid + 1 µM telmisartan internal standard) and gently mixed for 3 min. AS were analyzed in two ways: quantification and annotation of AS were performed using liquid chromatography coupled with quadrupole time-of-flight mass spectrometry (LC-QTOF-MS), and acyl chains were identified and quantified using gas chromatography coupled with mass spectrometry (GC-MS). Acylsugar extracts (4 µL) were run on an Ascentis Express C18 UHPLC column (100 mm × 2.1 mm, 2.7 µm) coupled to a Bruker Impact LC-QTOF-MS. The following chromatography conditions were used: column temperature 30°C, flow rate 0.3mL/min, Solvent A 10mM ammonium formate pH 2.8, Solvent B 100% acetonitrile, gradient: 0 min 95% Solvent A, 3 min 45% Solvent A, 13 min 0% Solvent A, 16 min 0% Solvent A, 16.01 min 95% Solvent A, 20 min 95% Solvent A. AS were annotated based on fragmentation patterns in ESI− modes. The following conditions were used: dry gas 10 liters/min, dry temperature 200°C, nebulizer gas 40 psi, and capillary voltage 2.8 kV. The mass range collected was *m/z* 50–1,500, and lock mass correction using sodium formate adducts [M-H]^−^ as the reference was applied during data collection. AS were manually annotated based on accurate mass and fragmentation patterns and quantified based on peak areas for extracted ion chromatograms of the [M-HCOO]^-^ adduct corrected for internal standard peak area and tissue dry weight (Schenck et al., 2022).

Acylsugar acyl chain analysis was performed from pooled acylsugar extracts using an Agilent 6890 GC-MS as previously described (Fan et al., 2020). Acyl chains were determined through library matches of the mass spectra of the corresponding ethyl esters using the NIST Library version 14. The relative abundances were calculated by integrating the corresponding peak area based on the total ion chromatogram (TIC) over the total acyl chain peak area.

### 6. Pest Resistance Evaluation

#### a. Bemisia tabaci assay

We first performed a no-choice bioassay using adult female *Bemisia tabaci* whiteflies placed in leaf discs (ø 12 mm) of *S. lycopersicum* cv. MT, MT-*Get*, MT-*Woolly,* and *Get02/Wo* (Suppl. Fig. 5). The leaf discs were placed adaxial side up, with the main vein inserted into 0.6% agar on the edge of vented Petri dishes. Ten female whiteflies were put on each Petri dish, and they were sealed and placed in an artificial climate box with controlled conditions (27°C, 16h photoperiod, 70% relative humidity) for 5 days. After 5 days, the number of whiteflies alive was quantified, and the number of eggs deposited on both foliar surfaces was counted.

An assay using clip cages was also performed. We used *S. galapagense*, MT, MT-*Wo*, MT- *Get*, MT-*Get02*, and *Get02/Wo* for this evaluation. Fifteen adult, healthy, female, and young whiteflies were placed in a clip cage on each plant. The plants were 5 weeks old, and the clip cages were placed on the 3^rd^ or 4^th^ leaf from the top. After 24 hours, the whiteflies were removed from the clip cages using CO2 sedation, and the eggs were counted.

#### b. Manduca sexta assay

Four-week-old MT-*Wo* and *Get02/Wo* plants were placed in a growth chamber (28°C, 11h photoperiod). Two newly emerged, healthy *M. sexta* caterpillars were placed on each plant. Two plants of the same genotype were placed together in an insect cage. The caterpillars were monitored daily to check survival and to replace the plant they were feeding on with a new one if needed. Seven days after the start of the experiment, each caterpillar was weighed.

#### c. Septoria lycopersici assay

In this assay, we employed a single spore isolate of S*. lycopersici*, sourced from naturally infected tomato plants in Long Island (New York). The fungal isolation entailed the pathogen extraction from infected tomato leaf tissues, then cultivation on V8 agar within 10-cm Petri dishes. After this, the plates were incubated at a controlled temperature of 23°C. After an incubation period of 10-12 days, conidia were meticulously harvested from the Petri dishes by inundating them with sterile distilled water. The inoculum concentration was adjusted to 5 × 105 conidia mL^-1^ using a hemocytometer. To ensure uniform spore deposition on tomato leaves during inoculation, a minute volume (approximately 10 μL) of Triton X-100 (2-[4-(2,4,4- trimethylpentan-2-yl)phenoxy]ethanol) was introduced into the inoculum suspension. This suspension was subsequently transferred to Nalgene aerosol spray bottles (ThermoFisher Scientific) for application. Artificial inoculation was carried out on 9-week-old tomato plants. These plants were positioned within a controlled humidity chamber, facilitating the maintenance of a relative humidity exceeding 95%, accomplished through a humidification system. Evaluating plant responses to the fungus involved a comprehensive examination of each inoculated plant for *Septoria* Leaf Spot (SLS) symptoms. This evaluation encompassed quantifying lesions (dark spots) on the 5^th^ oldest leaflets and determining the percentage of infected leaflets per plant. The resulting data was subsequently transformed into a *Septoria* tolerance score, graded on a scale ranging from 1 to 5, with the highest score of 5 signifying the highest resistance level.

## ACKNOWLEDGEMENTS

This work was supported by the USDA National Institute of Food and Agriculture (NIFA), Agriculture and Food Research Initiative (AFRI) (project # 2021-67014-33735). Part of the material used in this work was produced when E.V. was a pre-doctoral fellow at ESALQ/USP sponsored by the São Paulo Research Foundation (FAPESP – Brazil, grant # 2016/22323-4). We are grateful for the scholarship from the Council of International Programs of West Virginia (CIP WV) that allowed G.P.S. to work on this project. We thank “Conselho Nacional de Desenvolvimento Científico e Tecnológico” (Brazil) for the fellowship granted to L.E.P.P (CNPq grant # 308701/2022-4). Lloyd Sumner and Zhentian Li from the Metabolomics Center at the University of Missouri are acknowledged for their assistance with mass-spectrometry analysis, and Cassia Figueredo from the University of São Paulo for helping with scanning electronic microscopy.

## Author Contributions

EV, LEPP, and VAB conceived and designed the experiment, and wrote the manuscript; EV produced the tomato lines and performed the experiments; IOHL conducted the disease assay; RS, RT, PB conducted the herbivores assays; CLO and UR conducted the PACE analysis; TT and CAS conducted the metabolites profiling. GSP helped to collect and analyze the data; PB and CAS helped to revise the manuscript providing valuable suggestions. All authors read and approved the manuscript.

## Supplemental data

**Supplemental Figure S1.** Trichome quantification in the *Get02/Wo* recombinant population

**Supplemental Figure S2.** Chromatograms showing qualitative differences in AS profiles from the different genotypes.

**Supplemental Figure S3.** AS acyl chain composition from the different genotypes.

**Supplemental Figure S4.** Petri dish *Bemisia tabaci* no-choice bioassay.

**Supplemental Figure S5.** Comparative phenotypical analysis in the genotypes used.

**Supplemental Figure S6.** Complementary trichome characterization.

